# Monte Carlo Simulation of Diffusion MRI in geometries constructed from two-photon microscopy of human cortical grey matter

**DOI:** 10.1101/626945

**Authors:** Nima Gilani, Sven Hildebrand, Anna Schueth, Alard Roebroeck

## Abstract

**Purpose:** Neurodegenerative diseases such as Alzheimer’s disease cause changes and disruption to cortical microstructure and architecture. Diffusion MRI (dMRI) could potentially be sensitive to such changes. There is a growing interest in modeling of human cortical areas using a combination of quantitative MRI and 3D microscopy. The purpose of this study was to quantitatively characterize the cytoarchitecture of human cortical tissue from 3D fluorescence microscopy to simulate diffusion MRI (dMRI) signal in the cortex to better understand its diffusion signal characteristics.

**Methods:** Diffusion of water molecules and dMRI signal were simulated by an indirect geometry based method and a direct voxel based method in microstructural details extracted from microscopy of cortex. Additionally, residence times of diffusing spins inside voxel volumes were considered to set effective resolution limits. Mean diffusivity (MD) and kurtosis (MK) were calculated for variable cell and neurite densities, sizes and diffusion times under realistic values for permeability and free diffusion.

**Results:** Both simulation methods could efficiently and accurately simulate dMRI signals with fractional anisotropy, diffusion coefficient and kurtosis in agreement with previous reports. Simulated MD and MK showed changes with increasing diffusion times specific to cortical cell density and sizes, with MK showing the highest sensitivity. Intra-voxel residence times with increasing diffusion times showed that the effective dMRI resolution approaches the thickness of cortical layers.

**Conclusions:** Monte Carlo simulations based on 3D microscopy data enable estimating changes in MD and MK over diffusion times and are sensitive to cortical cytoarchitecture and its possible changes in neurodegenerative disease. When considering layer-specific cortical dMRI, effective resolution due to residence times is an important concern.

## Introduction

Changes in tissue microstructure of the cortex have been observed for diseases such as schizophrenia [1, 2], Alzheimer’s disease (AD) [3–5], and Huntington’s disease [4] in addition to age-related changes [6]. Diffusion MRI has been a successful and commonly used imaging modality to differentiate or characterize different diseases in the last decades [7] because of its sensitivity to tissue microstructure [8, 9].

It has been hypothesized that changes in cortical microstructure (e.g. cell density change or demyelination) result in observable variations in parameters derived from diffusion-weighted or diffusion-tensor imaging (e.g. fractional anisotropy, FA, or mean diffusivity, MD) [10–12]. For instance, Kroenke [10] discusses cases in which variations in FA have been of interest to study cortical development. Vrenken et al. [11] have shown decreases in FA of the cortex for patients with multiple sclerosis (MS). Microstructural changes associated with AD are decreased dendritic arborization, loss of synapses or dendritic spines [13–16] and selective loss of neurons [5, 17, 18]. Parker et al. have shown that these changes might result in decreases in orientation dispersion and neurite density indices (ODI/NDI) derived from neurite orientation dispersion and density imaging (NODDI) [12].

In addition, it has recently been shown that diffusion MRI can be used to characterise the architecture of cortical grey matter and its layers. Ganepola et al. [19] proposed the use of diffusion MRI derived parameters to redefine the widely used Brodmann’s areas [20] of the brain. Bastiani et al. showed delineation of cortical layer borders from high resolution dMRI on human post mortem tissue samples [21].

Human cortical grey matter has a dense composition (i.e. around 84% intra-cellular volume fraction) of axons, neurons, dendrites, glial cells, and blood vessels [22]. Despite this high level of cellularity, apparent diffusion coefficients measured from the cortex are relatively high compared to white matter. This could be explained by: a) the relatively large sizes of neurons (generally greater than 20 μm [2, 23, 24]) and b) glial minimum diameters of at least 2.5-3 μm [25], compared to very thin axons in white matter with diameters of generally less than 1 μm.

The layered composition of the cortex and partial volume effects at borders with cerebrospinal fluid (CSF) and WM, which have distinct diffusion characteristics, supposedly resulting in the diffusion signal to be multi-exponential. Initially, Maier et al. [26] proposed the use of biexponentials describing diffusion in cortical voxels adjacent to white matter or CSF. Biophysical multi-compartment models of diffusion, such as NODDI [27], can account for partial volume effects, and have recently been used to analyze diffusion in the cortex [28, 29].

Moreover, there are numerous reports of quantitative diffusion MRI of cortical grey matter in recent years [1, 11, 12, 26, 28, 30–32]. However, diffusion MRI of grey matter has received relatively less attention compared to white matter because of its smaller fractional anistropy and mylenation and hence less significant variations in diffusion MR related parameters.

Immunohistochemistry [5], scanning electron microscopy [33], different light microscopy techniques [34] such as two-photon microscopy [35], and ultra-high field (UHF) diffusion MRI [36] of *ex vivo* tissue samples have improved our understanding of the cortex microstructure in the last years. However, since there are a number of morphological changes in the fixation process [37–39] in addition to temperature differences, extrapolation from *ex vivo* MRI or microscopy to analysis *in vivo* diffusion MRI requires extensive mathematical modelling.

Here, we aim at simulating diffusion MRI signal in grey matter characterized by axon, neuron, dendrite, and glial composition, with an emphasis on cell-body composition, i.e. cytoarchitecture over different cortical layers. Monte Carlo simulations of diffusion as previously performed for white matter [40–42], and prostate [38, 43] were employed to study diffusion MRI signal in the cortex. We first consider residence times of water molecules in layered structures mimicking cortical grey matter boundaries to establish how much a voxel within a layer represents that physical layer without significant partial volume effects from other layers. Subsequently, we simulate diffusion MRI in cortical grey matter reconstructed from 3D microscopy using two different approaches: indirect geometrical reconstruction of cell bodies, and direct simulations on microscopy voxel grids. The simulations are performed for different microstructural compositions (density, size) of cell bodies and neurites, mimicking different cortical layers to simulate diffusion MRI signal in the cortex and to investigate its sensitivity to cortical cytoarchitecture.

## Materials and Methods

### Microstructural parameters derived from literature

There are numerous measurements of neuronal, glial, blood, or axonal cell sizes, cell densities or cell volume fractions in the healthy or diseased brain. The neural parenchyma is mostly made of neurons and glial cells and less than 3% its volume is made of the vasculature [44], hence most of these non-neuronal cells could be assumed to be glial cells [45]. Additionally, it has been shown that with increasing neuron diameters, glial/neuronal ratio increases; this has been hypothesized to be because of the increased metabolic needs of larger neurons [45, 46]. Accordingly, it has been hypothesized that glial/neuronal volume ratio might be of more interest to neuroscientists compared to glial/neuronal cell count [45]; this paradigm shift in cell measurements might benefit diffusion-weighted MRI analysis because diffusion parameters have been shown to highly rely on and have been described vs. cell volumes and volume fractions [38, 43, 47]. Fig. 1 is a scheme of the layered cyto and myelostructure of the cortical matter.

**Fig. 1.**
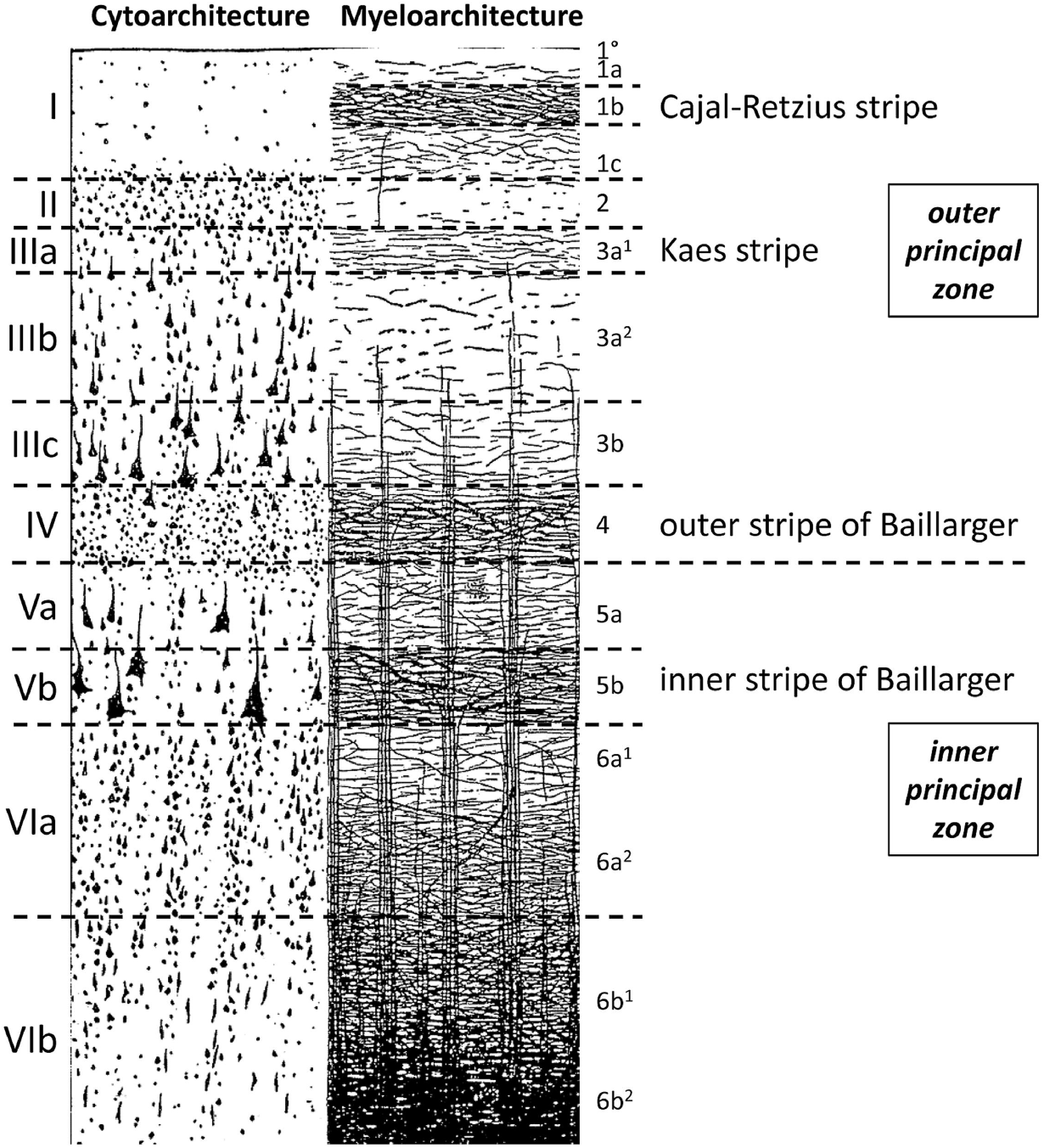
Layered myelo- and cytostructure of the cortical matter from [87, 88] under the terms of the Creative Commons Attribution License (CC BY, https://creativecommons.org/licenses/by/4.0/). Roman and Arabic numbers are for cytoarchitectonic and myeloachitectonic layers, respectively.

There are two main types of neurons in the cortex: pyramidal cells with diameters ranging from 20 to 120 μm, and granule cells which are star-shaped, have short axons, make local connections, and their diameters are around typically less than 20 μm [48]. There are a number of other studies giving similar estimates of neuronal diameters [2, 23–25, 49].

Average values of axonal diameters in the human cortical white matter have measured to be 0.5 to 1.34 μm (range of 0.19 to 4.79 μm) [50–52]. Deitcher et al. [53] have measured average dendritic lengths and diameters of 540-800 μm (dependent on depth from pia) and 0.76 ± 0.28 μm, respectively, in HL2/L3 of human neocortex.

The neuronal diameters are generally reported to be in the range of 11-27 μm (mostly around 20-25 μm) as discussed above. However, there are two issues that need to be addressed: first, pyramidal neurons have been assumed spherical in order to estimate their diameters. Second, the areas have been measured using two-dimensional images and such estimates of diameters are always an underestimation of cell sizes. Instead, there are a number of methods to recover 3D shapes of cells which are mentioned in [54]. Alternatively, high resolution and threedimensional two-photon microscopy acquisitions of the brain cortex could be employed to estimate neuron and glial cell sizes.

### Residence times of diffusion in cortical layers and imaging voxels

There are two different scenarios where the concept of residence time could help in feasibility assessment of separating physical diffusion compartments. The first case is having multiple microstructural compartments (each of them with their distinct diffusion characteristics) within each voxel; in this case, the low exchange between the compartments allows multiexponential fitting on the diffusion signal (e.g. [38] for the prostate, or [26, 27] for the brain). The second case is when the purpose is to check how much a voxel characterizing a microstructural layer is affected by nearby layers or neighbouring voxels. We take the second perspective here.

Here, residence times were measured for 3 different scenarios: one plate (mimicking the WM, GM boundary), two plates (mimicking cortical laminar structure), and six plates (mimicking a voxel).

#### One plate (single boundary)

Three-dimensional random walk simulation was performed in a geometry consisting of two compartments separated by an infinite plane (Fig. 2 a). The purpose was to show how long the water molecules at each initial position reside in the medium A dependent on the diffusion time andtheir initial position. The initial position of water molecules was normalized to 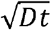 (where *D* was diffusion in the comparmtent and t was the acquisition time) and residence times were normalized to acquisition times. This normalization allowed for generalization of the simulation to any other similar geometry. However, here we interpreted the results for a specific case of white matter, and grey matter boundary.

**Fig. 2.**
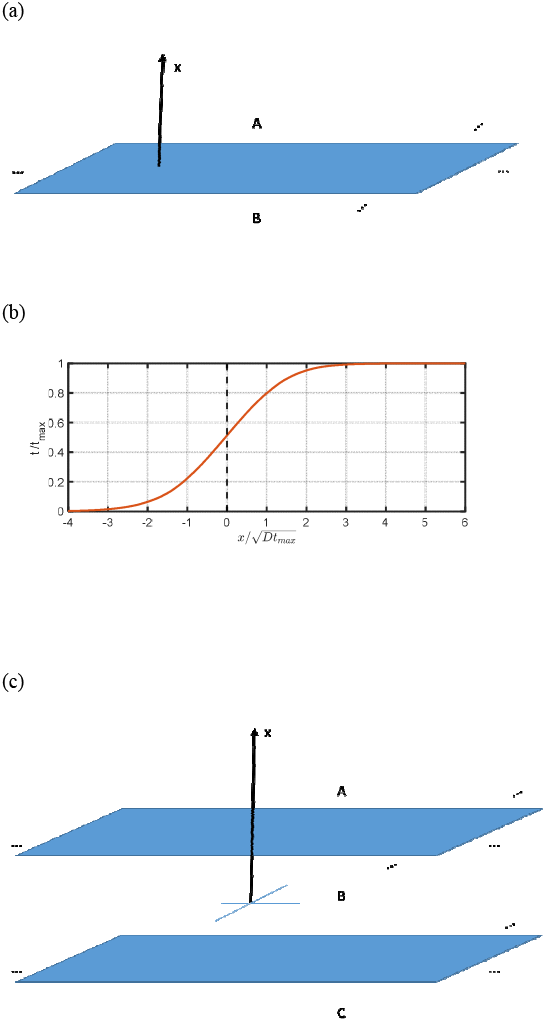
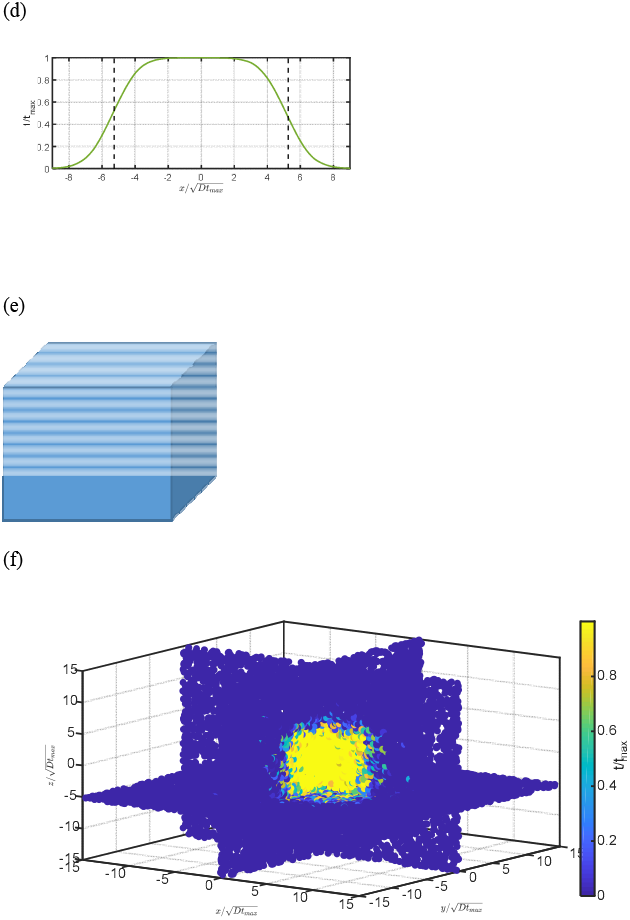
Residence time simulations in two mediums (WM and GM) separated by one plate (a) and (b), three mediums (cortical layers) separated by two plates (c) and (d), and inside a voxel (e) and (f). Notes: the dashed lines in (b) and (d) are x of the planes. For (b-f) all of the plates were either located at −5.27 or 5.27 /(*Dt*_max_)^0.5^

#### Two plates (cortical layer)

The three-dimensional random walk was performed in a geometry consisting of three compartments (medium) separated by two infinite planes (Fig. 2 c). The purpose was to determine residence time of water molecules in the central compartment relative to acquisition times for different initial positions. Later, results were normalized similar to the one plate simulation. While the results could be used for interpreting diffusion in any layered diffusion compartment, here the purpose of the simulation was to investigate residence times in the layered structure of the cortical matter.

#### Six plates (imaging voxel)

Similar to the one and two plate simulations, a simulation medium consisting of a voxel surrounded by 6 plates (Fig, 2 e) defining a cubic voxel was designed. Residence times of water molecules in the voxel were measured for different initial positions of spins in the voxel. This simulation gives a limit for the minimum voxel size for different acquisition times without the voxel being significantly affected by spins from the neighbouring voxels.

### Microscopy measurements

Realistic geometry of human cortical cytoarchitecture was reconstructed for simulation purposes using three-dimensional two-photon microscopy (TPM) on human cortical tissue. The tissue was optically cleared and fluorescently labelled for cell nuclei and cell bodies, using the procedures described in Hildebrand et al [55].

A filtering and Otsu thresholding based algorithm was developed to measure glial (*v_g_*), and neuron volume fractions (*v*_n_) using RGB images available of a Nissl-like fluorescent cell-body label [55]. Accordingly, the extracellular volume fraction (*v*_e_) was approximated as (1 – *V*_n_ – *V_g_*).

### Indirect geometry based simulation

A non-linear least squares based ellipsoid fitting function [56] was used in MATLAB Release 2017a (The MathWorks, Natick, Massachusetts) to characterize and estimate radii of neurons and glial cells separately. The function required at least 9 perimeter points to fit an ellipsoid; however, this condition was not adequate and an additional condition of having the points spread in at least 3 parallel planes was taken into account. Voxels corresponding to areas of glial nuclei and neurons in the visual cortex were segmented separately, and ellipsoids were fitted on them. Fig. 3 (a), and (b) are segmented areas of a neuron and a glial cell, respectively. *v*_n_, and *v_g_* ranges were 0.12-0.35, and 0.03-0.13, respectively, in 10 different TPM slides. The glial nuclei on average could be described as ellipsoids with the longer radii of 8.98±1.77 (ranging from 6.66 to 11.00) and smaller radii of 2.73±0.63 (ranging from 1.66 to 3.50). However, spheres better fitted on the neurons with average radii of 12.0±4.6 (ranging from 6.1 to 24.7) μm.

**Fig. 3.**
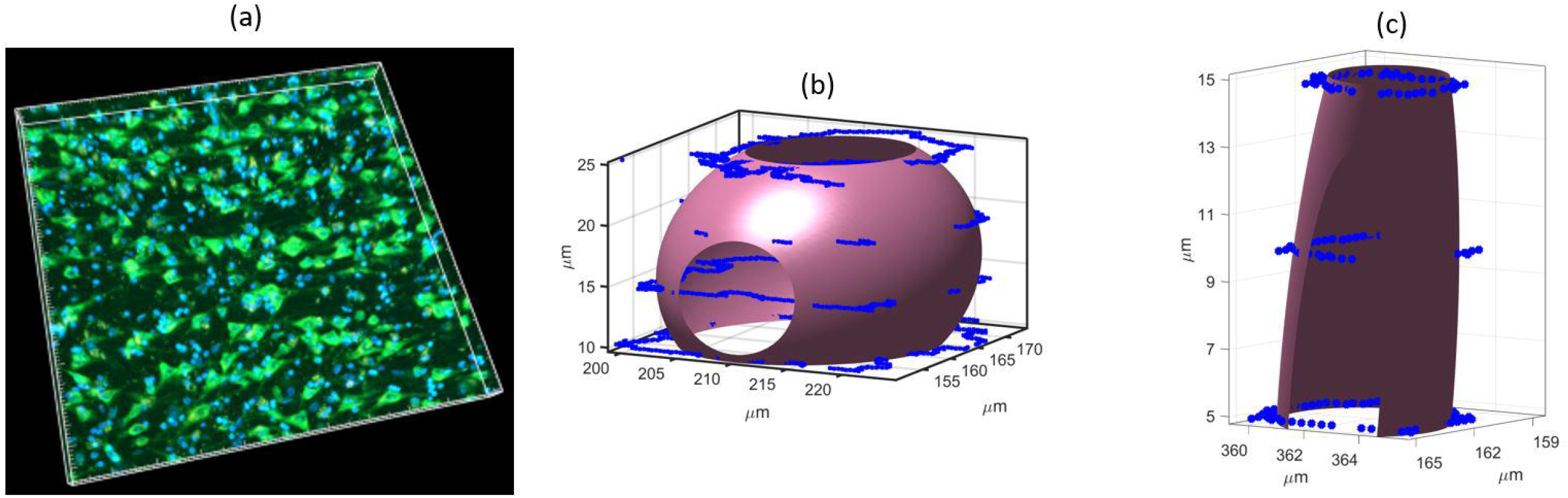
A 3D rendering of the 3D microscopy volume, with nucleus label in blue and cytoplasmic cell body label in green (a). Segmented areas of a neuron (b) and a glia (c) and the ellipsoids fitted to them. On average neurons were best described by spheres with radii of 12.0±4.6 and glia by ellipsoids with axes 2.73±0.63, 2.73±0.63, 8.98±1.77 μm in the visual cortex with N=50. Note: resolution and slice thickness of microscopy were 0.36×0.36 μm2, and 4.98 μm, respectively.

Mixtures of ellipsoids and cylinders with variable cell sizes and densities were used as geometry for simulation of diffusion in the cortex. Water molecules were randomly placed within the geometries and randomly moved with time steps of 1-25 μs corresponding to random walk steps of 0.09-0.48 μm assuming free diffusion coefficient of 1.6-1.7 μm^2^ms^-1^. Permeability of geometries was variable and simulated giving random chances of crossing to different permeability levels [38]. Pulsed gradient spin echo (PGSE) acquisition was simulated by recording changes in the phase of these randomly moving spins after the application of the diffusion gradients. Finally, diffusion kurtosis model [57, 58] was fitted on the signal. Detailed equations regarding the simulation of random walks, permeability, and PGSE could be found in [38, 43].

Neuronal radii of 10 or 20 μm were considered for all of the simulations. Glial cells were modelled as ellipsoids with radii of 2.7, 2.7, and 9 μm and three different radii of 0.3, 0.5, and 1 μm were considered for the axons. Different cortical areas or layers have different compositions; it is a time-consuming challenge to simulate and summarize all of these compositions in a single manuscript. Hence, here three different and simplified geometry scenarios which were most likely to simulate the visual cortex were considered.

### Simulation A (layer II/III)

The first scenario was the simulation of neurons and glial cells with varied neuron size and densities. Neuron volume fractions of around 0.1, 0.2, 0.3, and neuronal diameters of 10, and 20 μm were considered. This scenario mostly corresponds to layers II or III of the cortex.

### Simulation B (layer VI/white matter)

The second scenario was simulations of myelinated or unmyelinated axons with diameters of 0.6, 1, 2 μm and volume fractions of 0.1, 0.3, 0.6, 0.9; where for axonal volume fraction of 0. 9 hexagonal circle packing was employed and the other fractions were made by expanding the geometry phase space while keeping axonal morphology constant. It has been proven that hexagonal circle packing leads to the maximum possible volume fraction which is 0.9069 [59]. This simulation corresponds to layer six of the cortex, or in general white matter beneath the cortex. Permeability values of 10 and 30 μms^-1^ were considered corresponding to myelinated and unmyelinated axons, respectively.

### Simulation C (mixture)

The third scenario consisted of the mixtures of the two scenarios above. Neuronal and glial volume fractions were kept constant at 0.1, 0.05, respectively. Axonal volume fractions for this mixture varied between 0 to 0.3. This scenario mimics layers IV and V of the cortex.

For all of the simulations, 15 equally distanced *b*-values ranging between 0 to 2500 smm^-2^ were used. Four different diffusion times of 40, 60, 80, and 100 ms were used; the definition of diffusion times was similar to [37, 38, 43]. Two *D*_free_ (defined in [37, 38, 43]) values of 1. 6 and 1.7 μm^2^ms^-1^ were used for the simulations which have shown to be consistent with *in vivo* measurement of free diffusion which is affected by protein solutions [38] or consideration of an inherent *D*_free_ of 1.7 μm^2^ms^-1^ in NODDI algorithms [27, 28]. Additionally these values for *D*_free_ could also be verified by ADC estimates of 1.6-1.9 μm^2^ms^-1^ parallel to the direction of axons in reference [60], where these values are from a diverse range of 1.5-3 T Philips or Siemens scanners; note that diffusion parallel to the direction of fibre bundles in nearly free because axonal lengths are generally around or greater than a few hundreds of micrometres.

### Direct voxel based simulation

The ideal scenario for Monte Carlo simulation of diffusion is to directly use the geometry from microscopy in comparison with the mathematically reconstructed geometries of the previous section. This has been the subject of a recent study by Palombo et al. [61], where they have directly segmented area of cells from microscopy and simulated the random walks of thousands of spins within these geometries. However, the main issue with this approach is the long simulation times which will linearly increase with increasing of cell numbers, and the complexity of their area segmentation.

Additionally, a two-dimensional version of such direct simulation has been presented in [62] where only one cell type has been considered. However, they have not mentioned how threedimensional simulation mediums have been constructed from two-dimensional images (a recent reference [54] extensively discusses how this might be a formidable challenge).

Alternatively, here we used Matlab’s colour-based segmentation using K-means clustering (imsegkmeans) to separate different cell types by ascribing them different colors or numbers in two-photon microscopy samples. Microscopy voxels containing extracellular spaces were filled by zeroes, and other cell types were given a specific number (e.g. intracellular spaces of glial cells and neurons with respectively one and two).

Diffusion of water molecules was simulated within each microscopy voxel and if for each of the random walks the spin travelled to a neighbouring microscopy voxel with different geometry type, the available algorithm for permeability explained in the previous section was used to allow or reflect the random walk. Similar to [61], this method enabled different permeability ascriptions to each cell. Additionally, ascribing different numbers to different cell types allowed ascribing different permeability levels to cell wall of each cell type. Using this simplistic algorithm, random walk simulation of millions of spins took only a few hours on an Intel Xeon 3.60 GHz processor and there were not any requirements for complex reconstruction of biological geometries.

Accordingly, this method was applied to three-dimensional two-photon microscopy samples of the visual cortex previously reported in [55]. The samples had been labeled for nuclei (DNA) and cell-bodies (RNA). Glial cells tend to be rather small and dominated by their nucleus and are therefore mainly identified by the nucleus stain. Neurons on the other hand, tend to have large cell bodies and are therefore identified by both labels occuring in conjunction. Other acquisition details were: number of slices 21, slice thickness 4.98 μm, FOV 369.40×369.40 μm^2^, and matrix 1024×1024. Extracellular, neuronal and glial voxels were filled 0, 1, and 2, respectively. Both glial and neuronal cells were ascribed with permeability values of 30 μms^-1^ and random walk of water molecules was simulated in these samples.

## Results

### Indirect geometry based diffusion simulation

Figure 4 is a plot of D and K derived from the simulation of a few mixtures mixture of neurons, and glial cells mimicking layers II, and III of the visual cortex over varying diffusion times. The indirect geometry simulation method was used to reconstruct this geometry. For ADC, over diffusion times there is a near uniform ADC decrease for decreasing cell size and for increasing cell density (neuronal volume fraction, NVF). K better distinguishes cell sizes, especially for long diffusion times, where K increases with decreasing cell size. Over diffusion times there is a near uniform ADC decrease for decreasing cell size and for increasing cell density (neuronal volume fraction, NVF). K better distinguishes cell sizes, especially for long diffusion times, where K increases with decreasing cell size. Hence, higher diffusion times might help in better distinguishing neuronal cell sizes in pure grey matter voxels. However, kurtosis values are less variable with regards to neuronal volume fractions even if diffusion times are increased. Additionally, considering that apparent diffusion and kurtosis values are around 1-1.4 μm^2^ms^-1^ and 0.3-0.8,-2 respectively, the use of low *b*-values (generally less than 2000 smm^-2^) is recommended to decrease biases in estimating time-dependent diffusion and kurtosis. This is in agreement with earlier recommendations [63, 64].

**Fig. 4.**
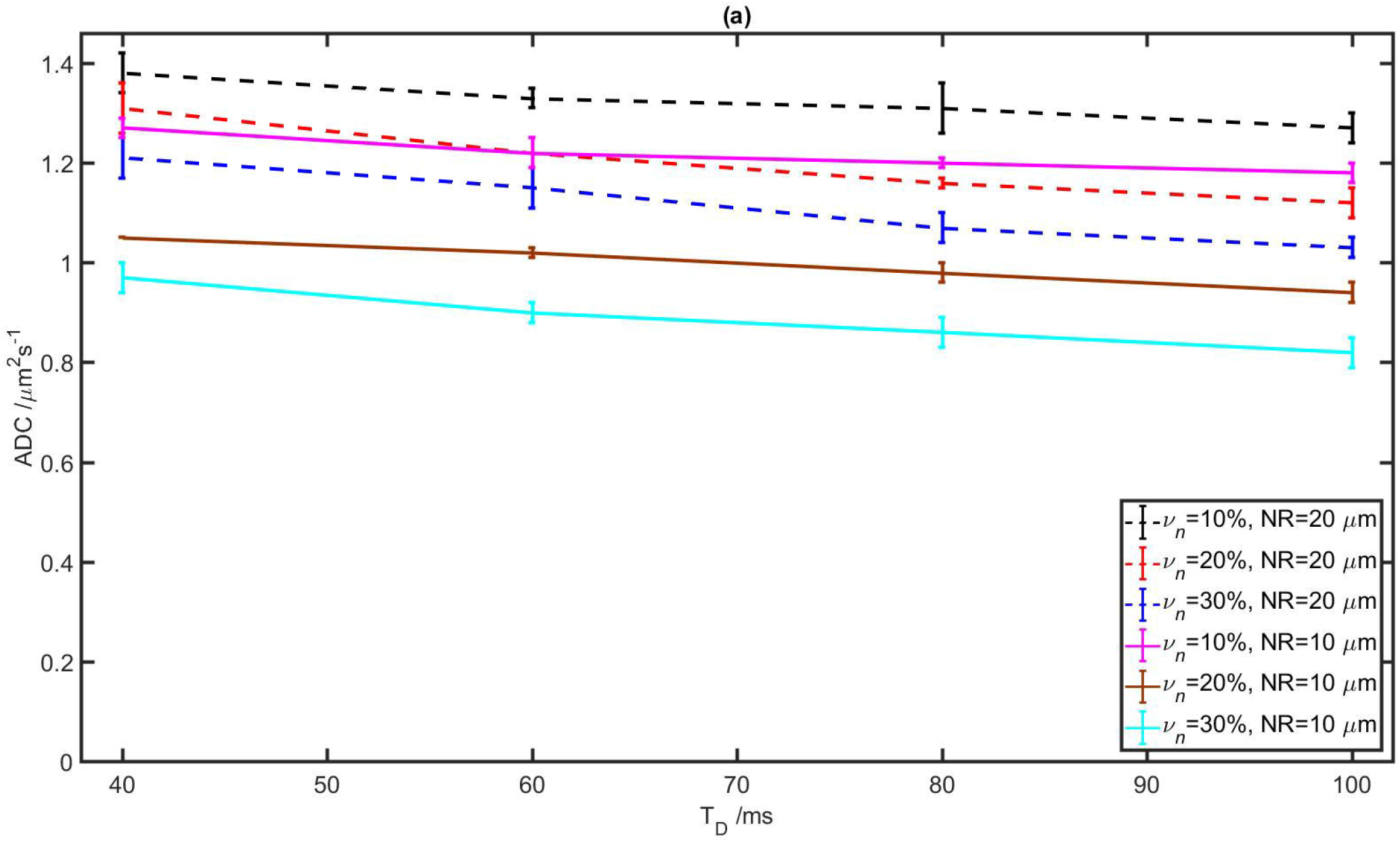
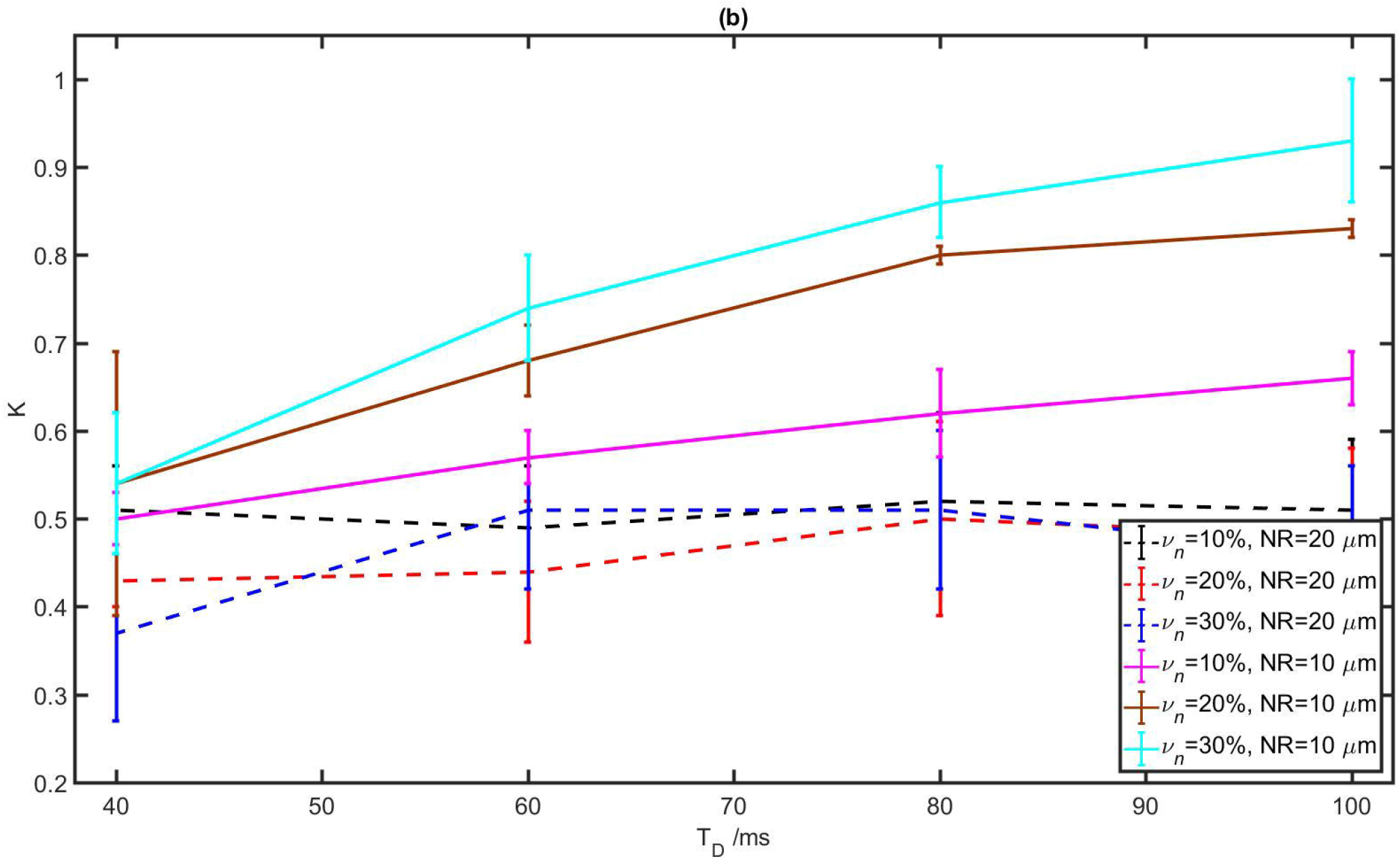
ADC (a) and kurtosis (b) of the simulated neuronal/glial mixture with variable neuron volume fractions (*v*_n_) and neuron radii (NR), for constant glial volume fraction, for diffusion times of 40-100 ms, with the indirect geometry based simulation method. Glial sizes were the same as Fig. 1 and their volume fraction was 0.03-0.07.

ADC of myelinated or unmyelinated axons with different cell sizes or volume fractions close to deep layers of the cortex are shown in Fig. 5. There are clear decreases in ADC with increasing myelination (decreasing permeability), and increasing axon radii. ADC curves for different axon radii and axon densities are almost parallel; this indicates that in the investigated, clinically feasible, ranges of acquisition parameters, variations in diffusion times do not substantially help in better characterization of axons. However, shorter diffusion times are slightly better in distinguishing of axons with typical small radii. In case of *ex vivo* acquisitions, there might be possibilities of using short diffusion times of less than 20 ms to improve axonal volume fractions and radii which has been indicated in [65, 66].

**Fig. 5.**
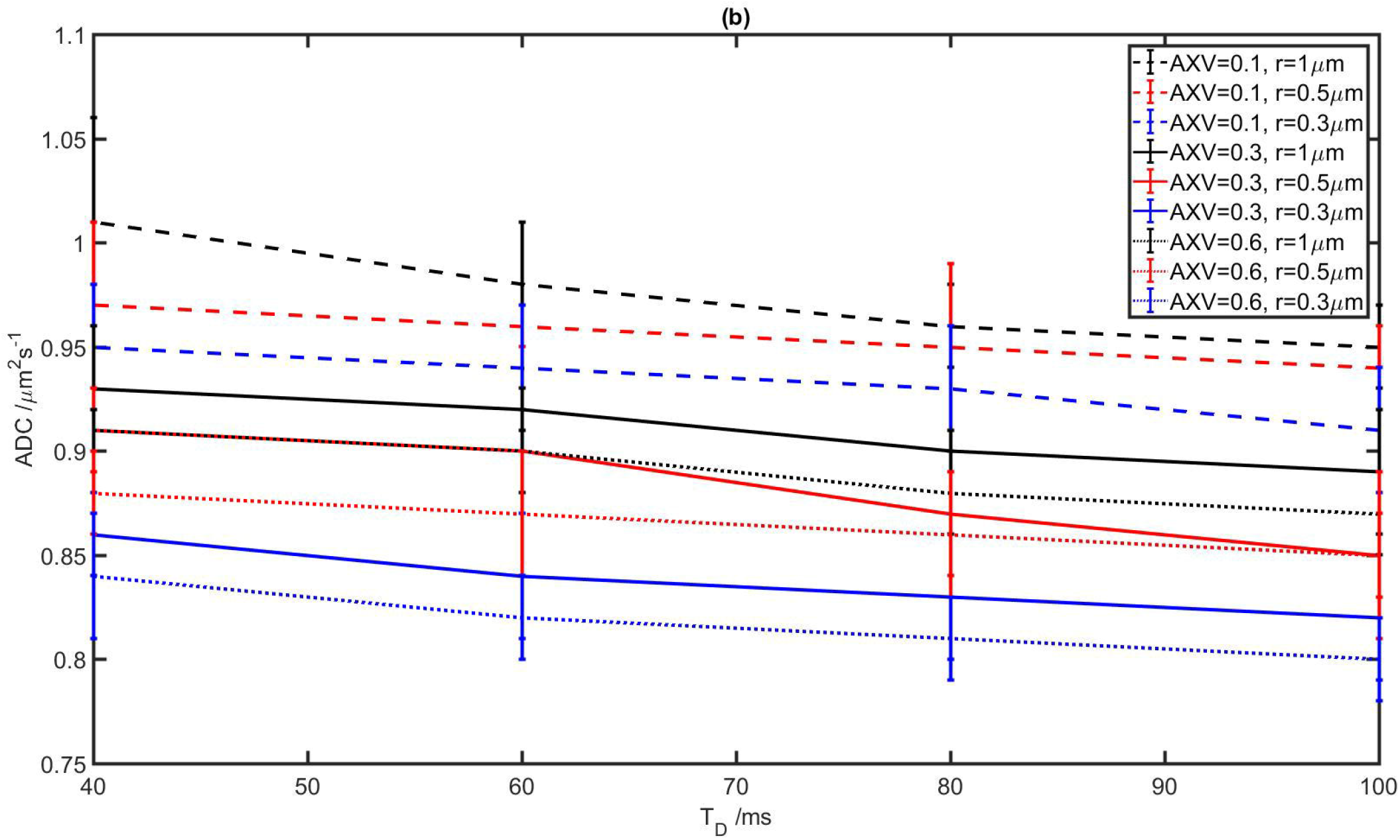
ADC simulated myelinated (a) and unmyelinated (b) axons with variable axonal volume fractions (AXV) and axon radii (r), in absence of cell bodies, for diffusion times of 40-100 ms, with the indirect geometry based simulation method.

It was observed that nearly all of the anisotropy in the cortex is caused by axons/dendrites and other cell structures such as neurons or glia have a negligible effect on anisotropy. However, this argument would not rule out that an anisotropic pattern of neuronal cells might have substantial impacts on the measured fractional anisotropy.

### Direct diffusion MRI simulation from microscopy

Fig. 6 is the measured ADC and K from the direct diffusion simulation in the two-photon microscopy sample. These values are in agreement with the simulations of Fig. 3 which could be approximately ascribed to geometries of layers II, and III.

**Fig. 6.**
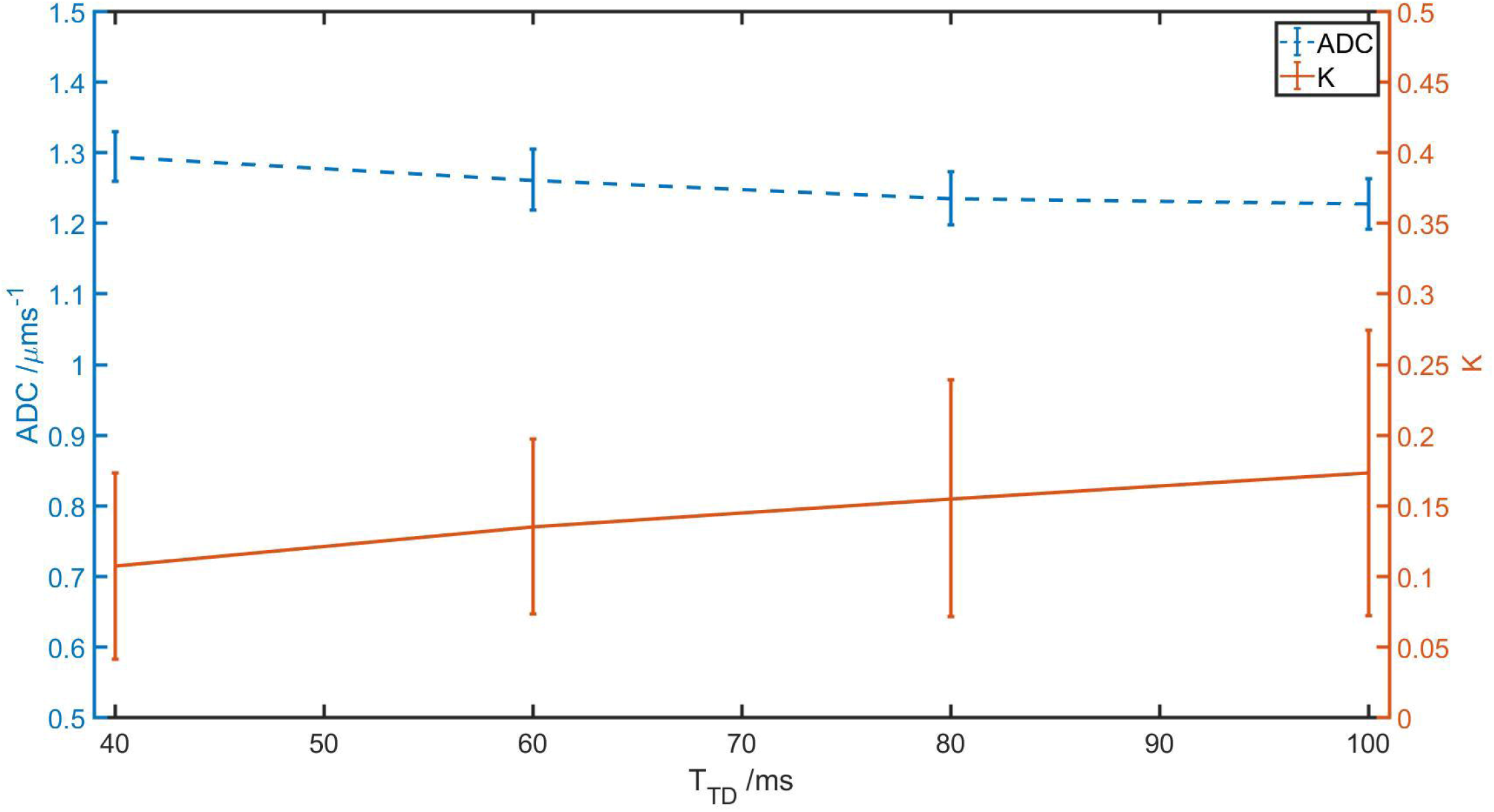
Direct simulations of (a) ADC and (b) *K* in a TPM microscopy sample of the visual cortex.

### Comparison with *in vivo* MRI

There are several reports of parameters derived from diffusion-weighted imaging in the cortical grey matter *in vivo,* some of which are summarized in table 1. These values are in a broad agreement with the simulations results.

**Table 1.**
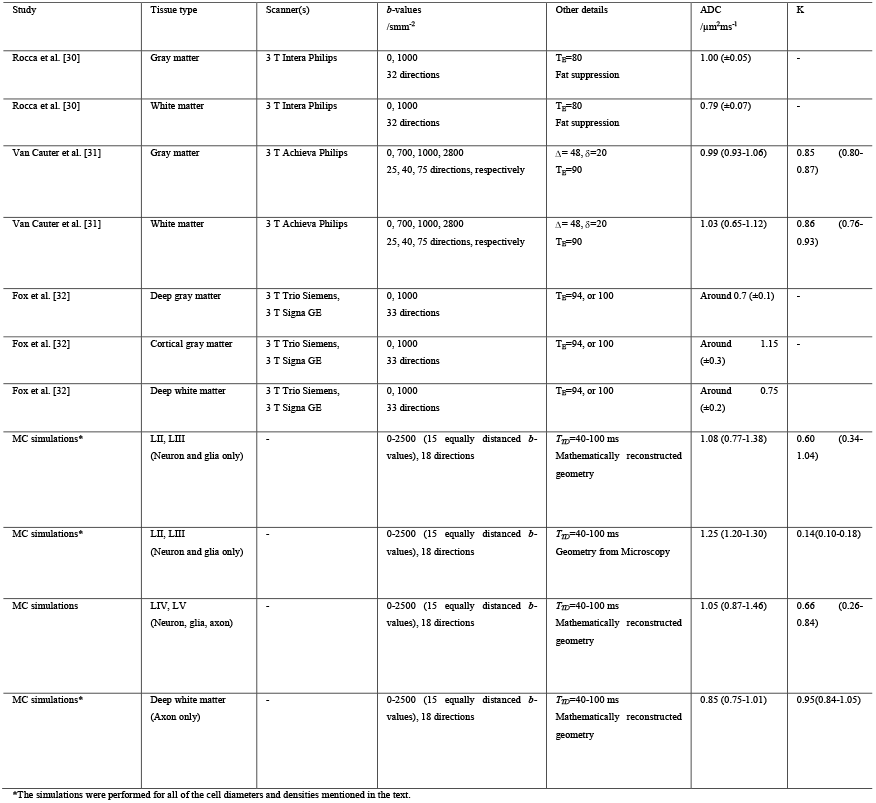
Diffusion-weighted measurements of the cortex from the literature.

### Residence time measurements

Figure 2 (b) gives the average residence time normalized to echo time for different starting positions from a single boundary with other boundaries being distant. This mimics the boundary of WM and GM or GM and CSF. Figure 2 (d) gives the average residence time normalized to echo time for different starting positions from two parallel boundaries. This mimics cortical layers where the boundaries are close. Figure 2 (f) gives the average residence time normalized to echo time for different starting positions in a 3D space surrounded by 6 plates (a voxel). It could be observed from the figures that with increasing diffusion coefficient in the tissue, and with increasing diffusion times water molecules reside less in their corresponding voxels or layers. In others words, water molecules near the boundaries keep bouncing back and forth in adjacent voxels or layers and the final MR signal is an average signal of them (partial volume effects).

Accordingly, decreasing diffusion times would help in better distinguishing of cortical layers. For example, assuming Fig. 2 (b) is representative of two adjacent layers with one of the layers having ADC of 1.6 μm^2^ms^-1^. Partial volume effects or average residence times could be calculated by averaging residence times of each compartment at different starting positions in figures 2 (b,d) if diffusion times (*t*_max_), layer thicknesses (dashed lines in Fig.’s 2 (b, d), and tissue diffusion coefficients (*D*) are given *a priori*. For example, with a diffusion time of 100ms, up to 14% of the signal from a 120 μm^3^ voxel at the boundary is from the adjacent layer. This layered partial volume effect would increase if longer diffusion times are used or if the layer is thinner. Additionally, there could be more partial volume effects for cortical layers (i.e. Fig. 2 (d)) where the same layer is adjacent to two layers, partial volume effects are around 30%. Additionally, these results could help in finding maximum feasible resolutions for each tissue type with regards to their characterisitc diffusion coefficient in addition to the discussions in [67].

## Discussion

McHugh et al. [67] have recently set resolution limits for diffusion weighted MRI. These limits are derived considering that mathematical estimates of tissue microstructure from signal in voxels with ranges of 1 mm are highly prone to noise. They have suggested lower resolutions and instead higher SNR values to better estimate diffusion parameters (specifically cell radii). Additionally, here we give resolution and diffusion time limits derived on the basis of high exchange or partial volume effects. Our limits would help in better characterization of cortical layers from diffusion MR *ex vivo.*

Fig. 4 is in agreement with previous studies [65, 66] stating that time-dependent diffusion measurements of axons with diameters of around 1 μm are sensitive to diffusion time at very short diffusion times (around a few milliseconds), whereas, for typically longer *in vivo* diffusion times of 40-100 ms changes in diffusion parameters vs. diffusion time are negligible. The same applies to simulations of grey matter [65]; however, at diffusion times of 40-60 ms more substantial variations in diffusion parameters relative to microstructural variations are observed compared to white matter. This might imply that considering the -s of clinical scanners, time-dependent diffusion measurements might more feasibly enable characterization of grey matter compared to white matter.

The results here further validate some of the commonly accepted arguments about correlations between microstructural changes and diffusion-weighted MRI observations. The examples are increases in permeability of axons (demyelination), increases in cell sizes (with constant cell density and structure), and increases in the heterogeneity of the mixture which result in decreases in fractional anisotropy, increases in ADC, and increases in kurtosis, respectively [43, 57, 58]. However, these simulations do not serve as proof but crossvalidation for such statements. Hence, care should always be taken when the aim is to estimate microstructural parameters from diffusion-weighted MRI [68].

It has been discussed in [69] that since there are different cell types in each voxel in addition to intrusions from CSF or WM, extrapolating the parameters derived from dMRI to variations in specific biophysical parameters is somehow more formidable compared to other tissue types such as the brain. For example, FA has a high correlation with myelination in white matter whereas this correlation is very weak for the cortex.

Jensen-Karger [57, 70] diffusion time-dependent kurtosis estimates could be used to derive rough estimates of MD and MK for two compartmental diffusion if diffusion times are long compared to residence times and also there is slow exchange between the compartments or permeability is low [38, 40]. Hence, it has been the trend of recent works by Lemberskiy et al. [71, 72] to keep the condition of long diffusion times (generally greater than 100 ms) in place and use stimulated echo acquisition mode (STEAM) diffusion MR and Jensen-Karger [57, 70] model to estimate time-dependent kurtosis. Alternatively, Monte Carlo simulations could be used to derive more degenerate and direct estimates of time-dependent kurtosis especially if the three required conditions for Jensen-Karger model are not met (i.e. there are more than two compartments, diffusion times are not long or there is high exchange). Monte Carlo simulations of Gilani et al. [38] is an example of bypassing the need to use numerous assumptions and equations [73] to estimate diffusion in a three compartmental tissue type such as the prostate when the diffusion times are not necessary long. Here, Fig. 4 (b) is one of similar Monte Carlo estimates of time-dependent kurtosis which interestingly is similar to the theoretical predictions of [57], however, this similarity could be by coincidence.

This study could also find implications for selecting optimized *b*-values in a simulation or in a real acquisition. Dependent on the purpose of MR acquisition (i.e. to estimate cells density cell type fractions), more specific *b*-value optimizations could be performed. Previously, Palombo et al. [74] have shown that the Stick model for diffusion is more sensitive to radii of soma at low *b*-value regimes (1-3 msμm^-2^), whereas, for a constant soma radius of 5 μm and ultra high *b*-values (3-15 msμm^-2^), the volume fraction of soma is more effective on the signal. The simulations presented here combined with optimizations of [63] also confirm their recommendation of using low *b*-value regimes to estimate large cell bodies. Whereas, for modelling axonal microstructure or estimating axonal radii, the use of higher *b*-values is generally recommended [41, 75, 76].

Diffusion MR has shown success in better characterization of the laminar mesostructure of the cortex in recent years [29]. Aggarwal et al. [77] have measured layer specific FA values *ex vivo* for Brodmann areas 4, 9, 17, and 18 and have plotted them versus cortical depths. These plots have relatively small inter-subject variability but are variable for each of these areas. FA values are generally less than 0.2-0.25 in the cortex and are substantially variable for each layer. In addition to the well known compositional effects on FA, they have also raised the possibility that decreases in FA of some layers could be caused by having tangential and radial crossing fibres in a voxel. Creating different perspectives for the cortex by different means such as the creation of gyral coordinate system [78] might also help in discovering more of neglected explanations for variations in diffusion signal.

We aimed here at quantifying the effects of variations in cortex composition and microstructure on changes in the diffusion signal. Simultaneous high-resolution MRI and light microscopy data will help in finding correlations between these; however, simulations could make them more specific. We did not confirm nor reject specificity of models such as NODDI or Stick model for diffusion. These models could be further investigated for the precision of model fitting either using Cramér-Rao lower bound such as in [79–81], finding the systematic biases in parameter estimations dependent on maximum *b*-values [63], missing analysis regarding how separate the compartments (if exchange rates are low) [43], or more in terms of how the models have been derived [82].

There were a few shortcomings for this work which might be alleviated in future work. First, simultaneous staining axons and neurons, glia, or in general all of the cortical matter microstructure would have been better for direct Monte Carlo simulations. While such staining is feasible for thin layers, it becomes challenging to thick layers of a few hundreds of micrometres which are more helpful for direct Monte Carlo simulations of diffusion.

Second, STEAM has recently been found applicable to such genre of studies *in vivo* or *ex vivo* [71, 83]. Simulating STEAM diffusion MRI is not inherently much more complex than PGSE and has been performed in [41]. Additionally, here we aimed at comparing the simulation results with abundantly available *in vivo* instead of *ex vivo* MRI data. This is because there are many differences between *ex vivo* and *in vivo* MRI such as fixation effects, temperature differences, etc. that makes direct extrapolation of *ex vivo* measurements to the analysis of in vivo MRI data a formidable task; this has been discussed for the prostate in [38] and for the brain in [39].

Simplified diffusion simulations in mathematically reconstructed geometries, or alternatively the novel direct simulations from microscopy might help in better microstructural characterization of diseases. Monte Carlo simulation of diffusion in these reconstructed geometries would help in improving our understanding of the physics of diffusion in each tissue type, and to better investigate existing models [82, 84]. Additionally, direct simulation of diffusion in different cortical areas if averaged for many samples would enable the creation of a library or dictionary [85] of direct diffusion simulation in the brain.

There might be a number of recent works with similar goals, this study however has some improvements such as successful simulation of permeability instead of assuming all micro-structural walls are impermeable compared to [86] or lack of requirement any complex mathematical and time-consuming construction of microstructure such as in [61] and instead direct simulation of microstructure from light microscopy data. Additionally, the work migh help in improving dMRI characterization of cortical layers (such as in [29]) or set additional resolution and diffusion time limits to better estimate microstructure from diffusion MRI from different perspectives following the work by McHugh et al. [67].

## Acknowledgements

NG and AR were supported by a Dutch science foundation (NWO) VIDI Grant (#14637). Additionally, AR was supported by an ERC Starting Grant (MULTICONNECT, #639938).

## Abbreviations

AD: Alzheimer’s disease
ADC: Apparent diffusion coefficient
AXV: Axonal volume fraction
CSF: Cerebrospinal fluids
Dfree: Free (unrestricted) diffusion coefficient
dMRI: Diffusion MRI
FA: Fractional anisotropy
GM: Grey matter
L: (Cortical) Layer
MD: Mean diffusivity
MK: Mean kurtosis
MS: Multiple sclerosis
NDI: Neurite density indice
NODDI: neurite orientation dispersion and density imaging
NR: Neuron radius
ODI: orientation dispersion indice
PGSE: Pulsed gradient spin echo
r: radius
STEAM: Stimulated echo acquisition mode
t: diffusoin time
tmax: diffusion (acquisition) time
TPM: Two-photon microscopy
UHF: Ultra high field
*V_e_*: Extracellular volume fraction
*v_g_*: Glial volume fractions
*v_n_*: Neuronal volume fractions
WM: White matter

